# Immune signaling mediates stromal changes to support epithelial reprogramming in Celiac duodenum

**DOI:** 10.1101/2025.03.05.641653

**Authors:** Dylan Richards, Klebea Sohn, Shrikanth Chomanahalli Basavarajappa, Daniel Horowitz, Joshua Rusbuldt, Lynn Tomsho, Kate Paget, Ivan Chavez, Koen Van den Berge, Brad McRae, Heather Loughnane, Seamus Hussey, Patrick Walsh, Darren Ruane

## Abstract

Coeliac Disease (CeD) is a chronic autoimmune disorder affecting 0.5-1% of the general population with a wide geographical distribution. Despite recent efforts to deeply phenotype gluten-specific immune activation at the single cell level, recent clinical studies targeting gluten degradation and other immune tolerance mechanisms have been unsuccessful. To this end, a deeper understanding of immune and non-immune cellular dynamics and interactions are required to characterize tissue-specific mechanisms responsible for CeD pathogenesis, repair, and resolution. Here, we assembled the most comprehensive scRNAseq dataset in Coeliac Disease to date, including 203,555 cells across 21 active CeD and 11 control duodenal samples. Compared to control duodenum, CeD was characterized by single cell differential changes in abundance, gene expression and cell-cell interactions across cellular compartments. In the immune compartments, CeD samples showed expected increases in plasma cell abundance and shifts toward type 1 effector biology (e.g., increase in cycling CD8^pos^, γδ T cells and IFNG transcriptional shifts) and T_fh_-related biology (e.g., increases in IL21 signaling to effector T cells). In addition, activated myeloid subsets, including DC2 and monocytes, were increased in disease and were characterized by increased pro-inflammatory pathway expression, including IL-1β. Non-immune compartments showed increased stem/crypt and secretory enterocytes in CeD samples with a decrease in absorptive enterocytes, reflecting the villus atrophy and crypt hyperplasia hallmarks of CeD epithelial dysfunction. Accompanying the epithelial changes, distinct changes in stromal populations were identified, particularly with increases in abundance and transcriptional activity of NRG1 and SMOC2 fibroblasts. Cell-cell interaction analysis across multiple cellular compartments proposed a distinct increased role of fibroblasts to support the epithelial reprogramming of the increased stem/crypt epithelial fraction in CeD, mediated by myeloid derived IL-1β signal and lymphoid-derived IFN-γ. This dataset reveals a previously unknown role for T-myeloid-stromal-epithelial cell communication in CeD, highlighting key mechanisms of the tissue-level cellular dynamics in response to gluten ingestion.

## Introduction

Coeliac Disease (CeD) is a common chronic autoimmune disorder with a wide geographical distribution, affecting 0.5-1% of the general population ^1^. CeD is characterized by a specific serological and histological profile associated with ingestion of gluten ^2^, in genetically predisposed individuals ^3, 4, 5^, resulting in inflammation of the small intestine. ^6, 7, 8^. The only treatment for the disease is a gluten-free diet (GFD) ^9^, generally leading to normalization of the gut mucosal pathology ^10^. Despite the strict requirements related to maintaining a GFD, a significant unmet clinical need exists in this population^11, 12, 13^, with most cases of CeD remaining undetected in the absence of serological screening and heterogeneous symptomology ^14^. Additionally, a proportion of CeD patients are refractory to a GFD, resulting in continual tissue damage and disease burden, impacting quality of life ^15, 16, 17, 18^.

The disease has a strong association with certain HLA-class II allotypes, including HLA-DQ2.5, HLA-DQ2.2 and HLA-DQ8. CeD has a strong hereditary component as confirmed by its high familial recurrence (∼10-15%) and high concordance of the disease among monozygotic twins (75-80%). Further, genome-wide association studies have identified more than 100 non-HLA related genes associated with CeD^19, 20^.

Seminal single-cell transcriptomics studies have enhanced our understanding of cellular dynamics associated with complex diseases within the gastrointestinal tract, including ulcerative colitis (UC), Crohns Disease (CD) and fibrostenotic CD ^21, 22, 23, 24^. These efforts have identified important cellular drivers of inflammation, remodeling, and tissue fibrosis, and as such, novel approaches may facilitate disease understanding in CeD pathophysiology. Various bulk transcriptomic and flow cytometric approaches have also enabled the interrogation of a select few lymphocyte populations In CeD ^25, 26, 27^, however, a comprehensive view of the immune and non-immune components of CeD in humans is lacking.

The pathophysiology associated with CeD disease is multifactorial, with a significant number of studies defining the role of T and B cell populations in mediating an aberrant adaptive immune response in disease^28, 29^. CD4^pos^ T cells of CeD patients are selectively activated by deamidated gluten peptides presented by disease associated HLAs. This occurs in the context of elevated IL-15 expression and facilitates the activation of intraepithelial lymphocytes (IELs) leading to small intestinal tissue damage highlighted by villous atrophy and crypt hyperplasia ^30^. These gluten specific CD4^pos^ T cells have been observed at high frequency in the Coeliac lesion of the duodenal bulb, with low frequency in the systemic circulation and are characterized by high expression of IL-21 and IFN-γ. Further, single cell RNA sequencing (scRNAseq) characterization of blood from CeD patients has highlighted an increase in gut homing (CD103^pos^), active (CD38^pos^) γδ and CD8^pos^ αβ T cells during gluten challenge. Moreover, a significant increase in B cells and the presence of anti-tissue transglutaminase 2 (TTG2) and anti-gluten peptide antibodies, have been reported in response to gluten challenge, confirming the importance of this T/B cell communication.

Despite a detailed understanding of the role of T and B cells in CeD pathogenesis, a significant knowledge gap remains regarding the complete immune and non-immune repertoire within the mucosa of patients with CeD. Key questions regarding the relationship between immune-stromal-epithelial cell compartments in active CeD remain undefined. Understanding this immune and non-immune cross talk, and the role of novel cellular subsets within the duodenal mucosa, will be key to defining novel therapeutics to support CeD patients and translate therapies into the clinic.

Our aim was to characterize the entire immune landscape associated with active CeD, using scRNAseq approaches, to elucidate the contribution of ill-defined CeD cellular subsets such as myeloid, stromal, and epithelial cells. Given the recent deconvolution of novel stromal and myeloid cellular subsets associated with Inflammatory Bowel Disease (IBD), we wished to identify and characterize tissue subsets and evaluate changes in specific cell types and populations. In tandem, our study aimed to uncover a deeper understanding of the cellular networks and receptor-ligand cross talk in the inflamed duodenal mucosa in CeD.

## Results

### A single-cell atlas of endoscopic tissue biopsies patients reveals a shift in cell type composition and cellular cross communication in Coeliac Disease

To generate a well powered scRNAseq transcriptional atlas, duodenal biopsies were isolated from 32 subjects, including 11 control and 21 active CeD (Fig. 1a). Post sampling, histological evaluation was completed to establish the relevant Marsh score for each subject, with representative histological images displaying the increasing villous atrophy and immune infiltration (Fig. 1b), a hall mark of CeD. All control subjects had Marsh scores of 0, while 19/21 active CeD had Marsh scores of 3a to 3c and 2/21 active CeD had a Marsh score of 2. Detailed sample metadata is available in supplementary Table 1. Biopsies for single cell analysis were cryopreserved prior to dissociation, Annexin^+^ de-enrichment was performed to enrich for viable cells and single cell sequencing processing. After data quality control and preprocessing, a total of 203,555 high-quality single cells remained for analysis, 136,317 cells from CeD and 67,238 cells from control.

**Figure 1:**
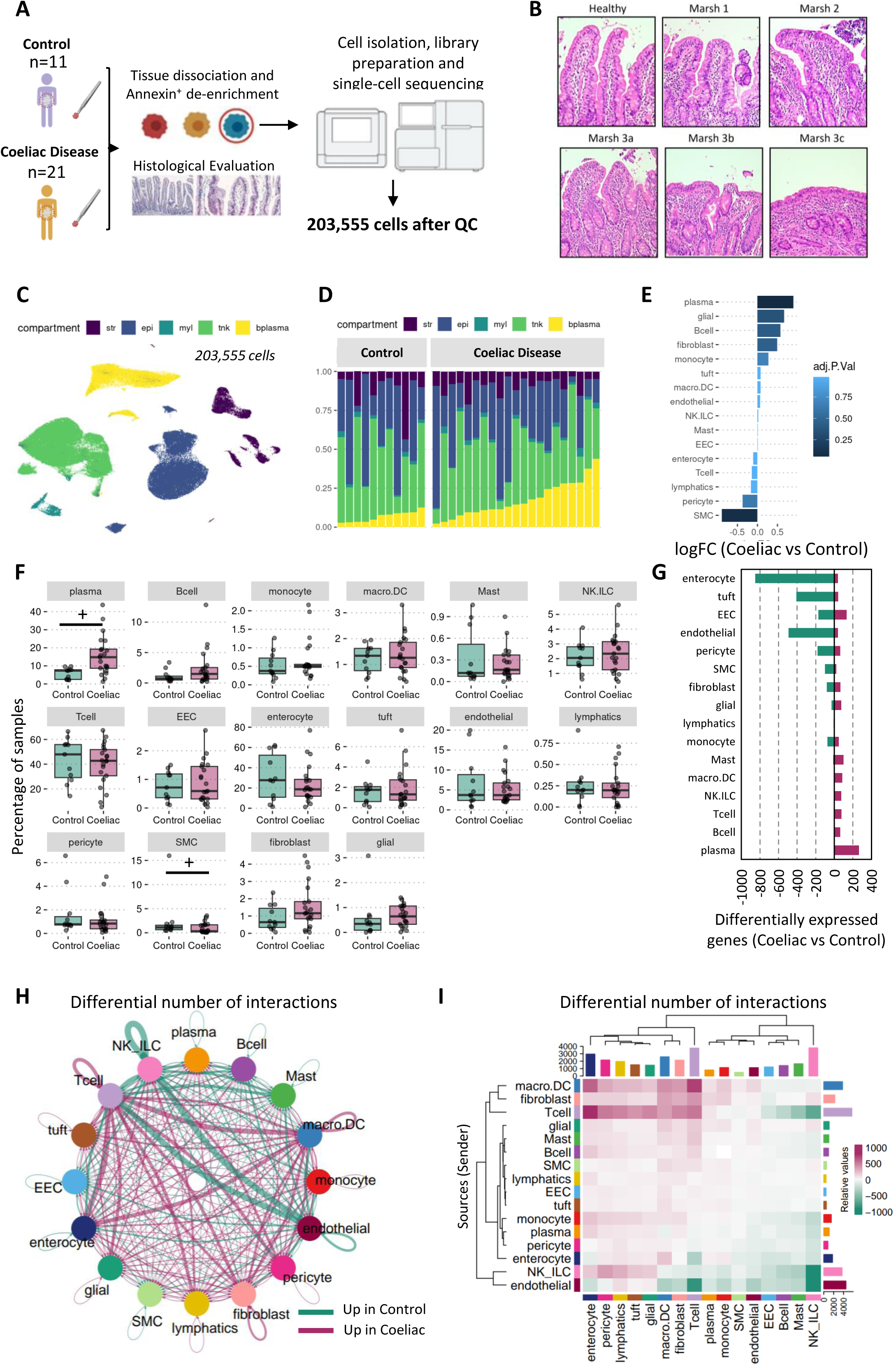
Single-cell atlas of duodenal biopsies reveals a shift in cell type composition, cellular gene expression and cell-cell cross communication in Coeliac Disease. A) Biospecimens from control and Coeliac subjects were collected for duodenum biopsy single cell RNA-sequencing and histological evaluation and for serum protein profiling. B) Representative H&E histologic images of each Marsh score indicating increasing immune infiltrate and villi atrophy with increasing severity. C, D) UMAP visualization and histogram of all cells collected, ordered by increasing plasma cell fraction, characteristic of Coeliac disease tissue. E) Differential abundance analysis of cell groupings indicate log-fold change (logFC) in Coeliac vs control samples, colored by adjusted P-value. F) Differential abundance visualized as percent of sample; + indicates adjusted p-value < 0.1 from differential abundance analysis. G) Differentially expressed (DE) gene counts (p < 0.05) per cell group generated from pseudobulk DE analysis. H, I) Cellchat analysis of cell-cell interactions with circle plot global view and heatmap visualization showing differential number of interactions between cell groups that are increased in Coeliac (red) or control (green).

The complete dataset was integrated across all subjects (using Harmony) and broadly annotated using canonical cell type markers to subcluster into stromal, epithelial, myeloid, T cell/NK and B cell/plasma compartments for further clustering and annotation per compartment (Fig. 1c). Characterization of the resulting cellular compartments revealed an overall dominant signal of increased plasma cell proportions in CeD samples, that was observed across Marsh scoring, consistent with existing literature in active CeD (Fig. 1d, Extended Fig. 1)^31^. Differential abundance analysis of cell types showed increases in plasma, B cell, monocyte, glial, and fibroblasts subsets in CeD samples (Fig. 1e,f). Interestingly, smooth muscle cells (SMC), were significantly decreased in CeD compared to control (Fig. 1f), which may reflect the atrophy of villi in CeD and associated nutrient (including lipid) malabsorption in CeD, as contractile villus SMC would normally support and propel dietary fat transport through the lacteal (i.e., adjacent lymphatic vessels) ^32, 33, 34^.

In addition to broad compositional changes related to active CeD, we examined the transcriptomic shifts in disease using pseudobulk differential gene expression analysis. Differential changes were observed across the tissue landscape with significant down-regulation of genes largely coming from epithelial and vascular cell types, while up-regulation of gene expression was observed across most cell types with the highest in immune cell types, reflecting the overall transcriptional impact of active CeD (Fig. 1g, Supplementary Table 2).

As a next step in understanding tissue-level shifts in transcriptomic activity, we examined the differential cell-cell interactions through CellChat receptor-ligand analysis. The total number of inferred interactions was increased in CeD, with 1550 additional interactions noted within coeliac samples, compared to control (Extended Fig. 2). Moreover, a significant number of receptor-ligand interactions were noted across various cellular subtypes in CeD. Notably, cellular crosstalk between myeloid and T cell subsets and a range of cell types, including fibroblasts and enterocytes, were significantly increased in the CeD mucosa. In contrast, crosstalk between endothelial cells and immune cell subsets such as ILC and T cells were differentially enhanced in the non-inflamed healthy mucosa from control patients (Fig. 1h,I).

In summary, differential changes in cellular composition, cell type, gene expression and cell-cell interactions characterize the tissue-level changes in CeD. To this end, a detailed compartmental analysis was conducted to enable further understanding of cellular crosstalk in the context of CeD.

### T_reg_, CD8^+^ cycling, γδ T and T_fh_-like CD4^+^ T cells are increased in the duodenum of Coeliac Patients

In the combined T/NK cell compartment (Fig. 2a) various T cell and Innate Lymphoid Cell (ILC) populations were identified, with an over representation of both CD4^pos^ and CD8 ^pos^ T cells. Populations of naïve and regulatory T cells were identified; these populations were characterized as CCR7^pos^SELL^pos^LEF1^pos^ and CD4^pos^FoxP3^pos^ respectively. A significant subset of CD4^pos^ T-follicular helper cells (T_fh_) were also identified as PDCD1^pos^ICOS^pos^. This population of T_fh_ also expressed IL21 and CXCL13, representing a cellular subset associated with CeD (Fig. 2b). Defined populations of NK NCAM1^pos^CD16^pos^, NK NCAM1^pos^CD16^neg^, and IL23R^pos^ RORC^pos^ KIT^pos^ ILC3-like innate lymphoid cells (ILCs) were also formally characterized (Fig. 2b). Further, diverse subsets of IELs with considerable heterogeneity in transcriptional states were also annotated based on ITGAE expression (CD103), compartmentalized into T-IEL CD8A^pos^, T-IEL TRDC^pos^, T-IEL CD8^neg^CD4^neg^, and cycling T cells expressing CD8A (Fig. 2b). The TRDC locus encodes the T cell receptor delta constant region, one component of the γδ T cell receptor, essential for the development of γδ T cells, which transcriptionally suggests a γδ T cell phenotype, known to share transcriptional qualities with CD8^pos^ T cells^25^. In addition, T cells expressing both TRDC^pos^TYROBP^pos^ phenotypes, with reduced CD8 expression, and a TRDC^pos^ LEF1^pos^ T cell with low ITGAE expression were noted across samples (Fig. 2b).

**Figure 2.**
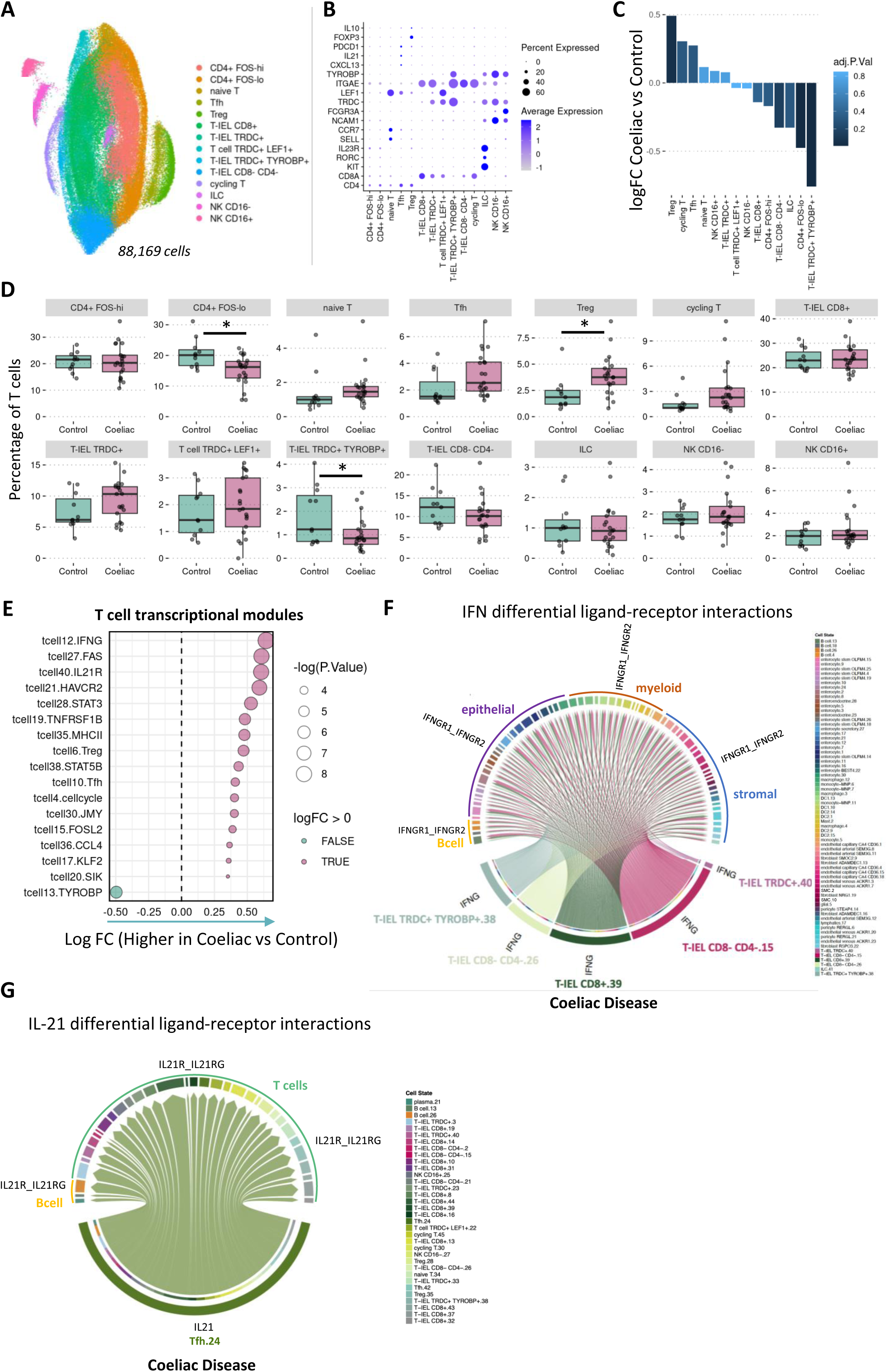
T_reg_, CD8^+^ cycling, γδ T and T_fh_-like CD4^+^ T cells are increased in the duodenum of Coeliac patients. A) UMAP visualization of TNK compartment (i.e., T cells, NK cells and ILCs). B) Expression of gene markers per cell type annotation. C) Differential abundance analysis of TNK compartment showing logFC of cell types in Coeliac vs control samples. D) Differential abundance visualized as percent of TNK compartment; * indicates adjusted p-value < 0.05 from TNK compartment differential abundance analysis. E) Differential gene module analysis performed using T cells (excluding NK and ILC cells) pseudobulk differential analysis, displaying modules with p-value < 0.05. F, G) CellChat chord diagrams highlighting ligand-receptor interactions of IFNG and IL21 signaling that are upregulated in Coeliac samples and absent in control samples.

Interestingly, various T/NK cell populations were found to be dysregulated between control and CeD subjects (Fig. 2c). An enrichment in T_regs_, cycling T, γδ T cells and T_fh_ cells as a proportion of the T cell compartment was observed in CeD disease (Fig. 2d). These cycling T cells, were characterized as CD8^pos^, expressed significant ITGAE and were found to increase from ∼0.8% to ∼2.5% of total T cells in active CeD. T_fh_ cells, characterized as CD4^pos^, PDCD1^pos^ and IL21^pos^, were also increased in CeD, likely reflecting gluten specific T cells, increasing from ∼0.8% to ∼3% of total T cells in disease. Furthermore, FoxP3^pos^ CD4^pos^ T_regs_ were also enriched within the duodenal tissue of CeD patients (Fig. 2b,d).

Coeliac subjects demonstrated an enrichment in TRDC^pos^ IEL T cells, likely reflecting an increase in γδ T cells in response to gluten exposure (Fig. 2b,d). In contrast, the proportion of IEL TRDC^+^TYROBP^+^ cells were markedly reduced in the mucosa in CeD (Fig. 2b,d). As TYROBP (DAP12) has been reported to be exclusively expressed by natural IELs, this significant decrease is consistent with previous observations and indicates a loss of a subset of γδ T cells in CeD, possibly as a consequence of disrupted homeostasis and reduced mature intestinal epithelial cells post gluten exposure in CeD ^25, 35^.

No significant changes in cellular frequency were noted in ILCs (IL23R^pos^ RORC^pos^ KIT^pos^) or NK NCAM^pos^ CD16^pos^ or NK NCAM^pos^CD16^neg^ cells (Fig. 2d). CD4^pos^FOS^low^ T cells were significantly reduced in CeD, suggesting a reduction CD4^pos^ quiescent T cells, given the exposure of CeD patients in our study to a gluten rich diet, a reduction in quiescent T cells and shift to an activated phenotype is expected.

To contextualize the transcriptomics changes more fully in the CeD mucosa, with regards to transcriptional state shifts within the T cell compartment, and to account for co-expression patterns between genes, we further examined T cells leveraging differential module analysis. To derive disease-relevant, cell type-specific transcriptional state modules, briefly, gene correlation networks were constructed and clustered for each cellular compartment of interest (e.g., T cells). Modules were then used with gene set variation analysis (GSVA) to generate module scores for pseudobulk cellular samples for differential module analysis. Module lists characterizing processes of shared co-expression across a cell group can be found in Supplementary Table 3. Differential module analysis of T cells found a significant upregulation in T cell transcriptional states associated with IFN-γ and type 1 biology in CeD: the IFN-γ module (tnk12.IFNG) incorporates a variety of genes including IFNG and GZMB, reflective of an increase in effector Type 1-like transcriptional processes in CeD, a key feature of CeD immune landscape. Significant increases in T cell modules defined by co-expression with FAS, HAVCR2 (i.e., TIM3), and CCL4 were also observed in Coeliac disease biopsies (Fig. 2e) as were IL21R and TNFRSF1B (TNFR1), all associated with an activated inflammatory profile. In addition, consistent with an enrichment of T_regs_, cycling T and T_fh_ cells, significant increases in the regulatory T cell transcriptional modules (including genes such as FOXP3, BATF), cell cycle module, and T_fh_ modules (including genes such as CXCL13, CXCL21) were also observed (Fig. 2e).

To examine the downstream effects of the transcriptional shifts in CeD T cells, CellChat differential ligand-receptor analysis was performed and further highlighted two key upregulated ligands: IFNG and IL21. A significant enrichment in IFN-γ inferred interactions from IEL T cell populations (including T-IEL TRDC^pos^, T-IEL CD8^pos^, and T-IEL CD8^neg^CD4^neg^ cells) to many cell types with IFNG receptors across myeloid, epithelial, stromal, and B cell compartments (Fig. 2f). These results serve to contextualize the broader cellular crosstalk evidence in the CeD duodenal mucosa for the orchestrating role of IFN-γ from gluten-induced cytotoxic T cells as previously reported ^25^. Furthermore, a significant increase in IL-21 signaling from T_fh_ cells to various T cell and B cell populations was also identified (Fig. 2g). This, along with the increase in T_fh_ transcriptional processes, as well as IL21R, STAT3, and STAT5B co-expression modules, support the inferred signaling of IL-21 within the CeD T cell compartment.

In summary, the T cell compartment is significantly dysregulated in CeD disease, as characterized by an increase in T_regs_, cycling CD8^+^ T, γδ T and CD4^+^ T_fh_ like cells. In addition, significant increases in IFN-γ signaling in IEL T cells in addition to IL21 signaling mediated by T_fh_ cells, represents a significant contribution to local tissue immune responses, remodeling, and disease pathogenesis. Furthermore, the IFN-signaling from activated T cells in CeD has the capacity to impact immune and non-immune relevant cell type in the duodenal tissue including myeloid, epithelial, and stromal cellular subsets.

### Activated myeloid cellular subsets are enriched in Coeliac Disease mucosa

Unique to our dataset is the comprehensive characterization of other immune and non-immune subsets in CeD, beyond T and B cells. These included mononuclear phagocytes (MNPs), comprising DC and monocyte subsets with the former characterized by a gene program that includes the DC hallmark receptor FLT3, while the latter expressed a variety of core macrophages genes including ZEB2, a transcriptional factor required for maintaining the tissue-specific identity of macrophages.

In the DC, myeloid and mast cell UMAP, high resolution characterization revealed a variety of cellular subsets including DC1, DC2, macrophages, monocyte MNP, monocytes and mast cells (Fig. 3a). These cellular subsets were characterized and differentiated based on known transcriptional markers (Fig. 3b). Detailed transcriptional evaluation of these subsets identified DC1^+^ cells, as CLEC9A^pos^FLT3^pos^ZEB1^pos^ (Fig. 3c). DC2^+^ were characterized as CD1C^pos^ID2^pos^ITGAX^pos^CD68^pos^ (Fig. 3c) while monocytes and macrophages were identified as CD14^pos^CD68^pos^ZEB2^pos^ and CD68^pos^CD163^pos^FTL^pos^ respectively (Fig. 3c). These macrophages were broadly characterized by a lack of pro-inflammatory gene expression, while both subsets expressed significant MRC1 (CD206) levels, a marker of gut resident macrophages. Interestingly, various myeloid subsets including monocytes, DC1 and DC2 expressed significant levels of IFNGR1 and IFNGR2 (Fig. 3c), endowing these cells with heightened IFN-γ signaling/responsiveness. In this context it is noteworthy that previously discussed IEL T cell analysis, as characterized by cell chat in Fig. 2f, was suggestive of significant cellular crosstalk between activated gluten specific CD8^pos^ and MNP cellular subtypes in CeD, potentially modulating their phenotype profiles, in the context of gluten exposure.

**Figure 3.**
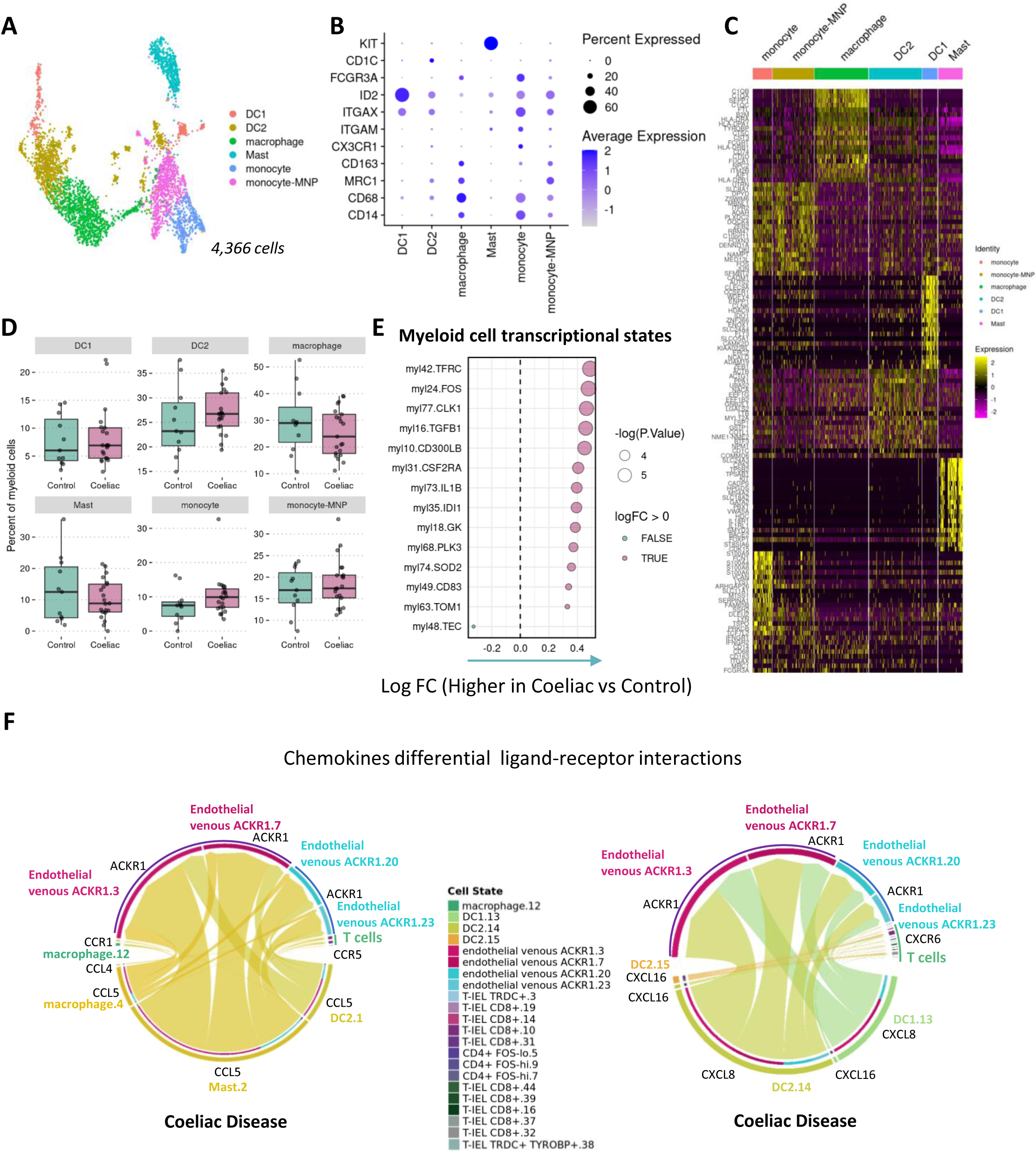
Activated myeloid cellular subsets are enriched in Coeliac Disease mucosa. A) UMAP visualization of myeloid compartment. B) Expression of gene markers per cell type annotation. C) Detailed heatmap highlighting genes that characterize myeloid cell types in duodenal biopsies. D) Boxplots of myeloid subtypes as percentage of myeloid compartment. E) Differential gene module analysis performed using myeloid pseudobulk differential analysis, displaying modules with p-value < 0.05. F) CellChat chord diagrams highlighting chemokine ligand-receptor interactions from myeloid cells that are upregulated in Coeliac samples and absent in control samples.

Cellular abundance analysis revealed distinct changes within the myeloid compartment in CeD biopsies. Relative to the total sample, monocyte cell group (i.e., monocyte and monocyte-MNP) increased in CeD, suggesting a modest increase in CD16^pos^ infiltrating cells (Fig. 1e). Relative to the myeloid compartment, conventional DC1^+^, DC2^+^ cells, and monocytes showed relative increases in CeD, whereas mast and macrophages showed relative decreases in CeD (Fig. 3d). Beyond the relative shifts in cell numbers, detailed differential module analysis (Fig. 3e) revealed an increase in inflammatory processes across various myeloid cellular subsets in CeD. Myeloid subsets had increased transferrin receptor (TFRC) pathway activation in CeD. The TFRC pathway is associated with apoptosis and cellular activation in addition to mediating iron uptake, with iron deficiency a key characteristic noted in patients with CeD^36^. Significant increases in the IL1β module (including NLRP3, FPR1) and SOD2 module (including CXLC2) were also observed in myeloid cells derived from CeD samples, reflecting a pro-inflammatory cascade signaling outward in response to gluten exposure. Further, enhanced FOS, CLK1 and CD300LB signaling also characterizes the myeloid state in CeD, suggestive of broad cellular activation, including RIPK2 and TNFRSF10D. Increases in the TGFβ1 module (including ADAMTSL4, CREB1 and FOSL2) within the CeD myeloid compartment indicates myeloid cells as a source for stromal interactions and for TGFB1-induced epithelial regeneration (Fig. 3e). These data highlight the significance of an activated myeloid population contributing to CeD pathogenesis and tissue remodeling upon gluten exposure.

The contribution of this activated myeloid phenotype to CeD pathogenesis and tissue remodeling was also confirmed through cell-chat analysis. Here, activated myeloid cells including DC1, DC2, mast cells and macrophages produced a variety of chemokines-including CCL4 and CCL5 acting on downstream CCR1, ACKR1 and CCR5 expressed on epithelial, stromal and T cell subsets (Fig. 3f). DC1 and DC2 subset expression of CXCL8 and CXCL16 was also increased in CeD. CXCL8 and CXCL16 ligand receptor interactions from DCs, included epithelial, stromal and T cell subsets, suggestive of a key role for DCs in mediating cellular recruitment and interaction with these cell types in the context of CeD.

These data reflect a dynamic myeloid activation state associated with active CeD and reveals significant cellular crosstalk between T cells, myeloid and downstream stromal/endothelial cells in CeD, previously undescribed.

### Intestinal epithelial cellular dysfunction and enhanced secretory enterocytes are a hallmark of Coeliac Disease

In addition to the immune compartment, a variety of non-immune subsets were also characterized, with the aim of evaluating their contribution in CeD and to assess cellular communication dynamics within the duodenal mucosa. With our cellular preparation we were able to identify three distinct enterocyte subsets, representing stem, secretory, and absorptive lineages (Fig. 4a). Stem-like enterocytes were broadly characterized as OLFM4^pos^ (i.e., enterocyte stem OLFM4 population) and included LGR5-expressing cells, reflecting the crypt stem cell phenotype, and MKI67-expressing cells, indicating a transit amplifying population present. Absorptive enterocytes included the presence of BEST4^pos^MEP1A^POS^ (enterocyte BEST4) and a putative mature population of MEP1A^POS^GSTA1^POS^CA2^POS^ cells (enterocytes). Secretory enterocytes encompassed tuft cells (expressing TRMP5, DCLK1), enteroendocrine cells (expressing range of gut hormone genes like CHGA and PYY), and goblet-like cells (annotated “enterocyte secretory”) that expressed a range of mucosa-supporting genes like MUC1, MUC5AC, TFF1-3, and REG4 (Fig. 4b). Expression of MUC5A within the secretory enterocytes may reflect cellular subsets of the Brunners glands. Despite our ability to recover many crypt-like stem cells with our protocol, limited expression of markers for the crypt small intestine Paneth cell, such as DEFA5 and DEFA6, were observed within our dataset, suggesting a epithelial-specific processing step may be required to adequately sample this limited cell type.

**Figure 4.**
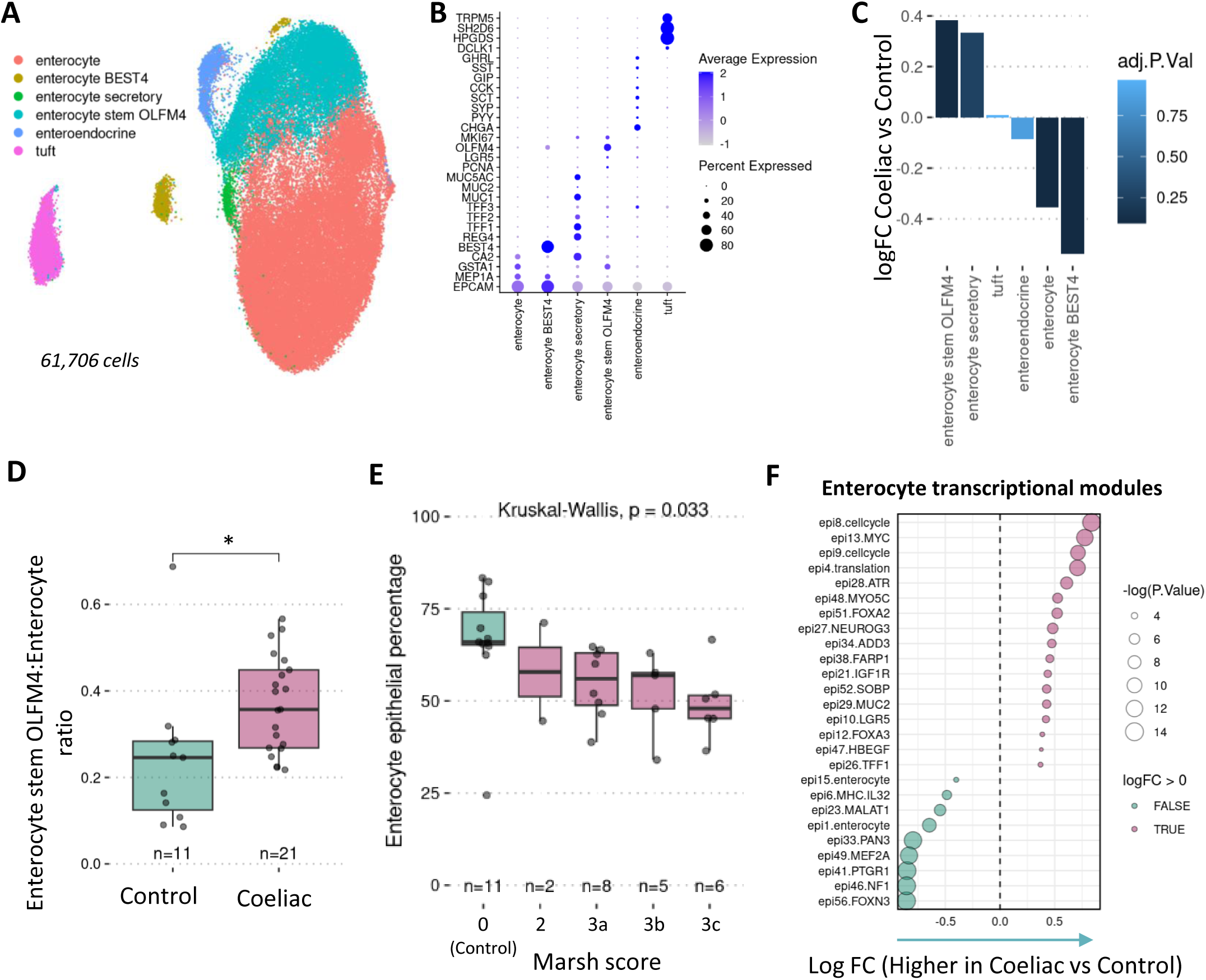
Intestinal epithelial cellular dysfunction and enhanced secretory enterocytes are a hallmark of Coeliac Disease. A) UMAP visualization of epithelial compartment. B) Expression of gene markers per cell type annotation. C) Differential abundance analysis of epithelial compartment showing logFC of cell types in Coeliac vs control samples. D) Ratio of enterocyte stem OLFM4 cells to enterocyte counts per sample; * indicates P-value < 0.05 from independent T-test. E) Enterocyte epithelial percentage, as a function of clinical Marsh scoring; P-value marked from Kruskal-Wallis test. F) Differential gene module analysis performed using enterocyte pseudobulk (i.e., enterocyte, enterocyte stem OLFM4, enterocyte secretory, enterocyte BEST4) differential analysis, displaying modules with adjusted p-value < 0.05.

Within the duodenal tissue of CeD, a significant shift in enterocyte phenotype and frequency was noted. Relative to the epithelial compartment, increased abundance of stem-like enterocytes (OLFM4) was observed in disease, in addition to significant increases in secretory enterocytes (Fig. 4c). In contrast, the absorptive populations, BEST4 enterocyte and enterocyte cells, were significantly reduced. These cellular shifts are further highlighted by a significant increase in stem:mature enterocyte ratio in CeD (Fig. 4d). Marsh histological grading accounts for increasing levels of villous atrophy within grade 3 and is reflected in CeD samples by significant decreases in enterocyte epithelial percentage across Marsh scoring in CeD (Fig. 4e).

Differential transcriptional module analysis supported the associated processes observed in the changing epithelial compartment. In CeD, enterocyte transcriptional module analysis revealed an increase in putative crypt phenotypes (Fig. 4f) as characterized by the up-regulated cell cycle module and LGR5 (including OLFM4) transcriptional modules. Upregulation of the MUC2 and NEUROG3 modules in enterocytes from CeD patients also reflects increasing secretory enterocytes and processes to support epithelial differentiation biology, as NEUROG3 has been reported to regulate differentiation of secretory cells ^37, 38^. Consistent with a reduction in enterocyte frequency alongside increases in stem/secretory enterocytes, a reduction in mature transcriptional processes (e.g., epi1 EPCAM/MEP1A-containing module) was also observed (Fig. 4f).

In total, the overall shift toward reduced absorptive enterocytes and increased crypt/stem activity reflects the villous atrophy and crypt hyperplasia hallmark phenotypes, respectively, observed in CeD epithelium^30^.

### Coeliac Disease is characterized by unique tissue relevant stromal cellular subsets reflective of disease re-modeling and repair

Recent studies in RA, UC and CD have reported distinct disturbance in the stromal cellular compartment associated with disease pathogenesis. However, little is known about the composition of potential small intestinal stromal cells towards mucosal inflammation in CeD. To evaluate the stromal cellular landscape in CeD, we performed detailed cellular deconvolution to identify novel duodenal stromal subpopulations. In the combined stromal compartment, we identified a total of eleven distinct cellular subsets, primarily grouped into glial, fibroblast, endothelial and SMC populations (Fig. 5a). Endothelial subsets characterized as either ACKR1^pos^, CD36^pos^ or SEM3G^pos^ representing venous, capillary, and arterial vasculature, respectively, as well as lymphatic endothelial cells were also identified in duodenal stroma with minimal changes in abundance (Fig. 5b,c). Pericytes, similarly characterized in IBD ^24^, were identified as STEAP4^pos^ or RERGL^pos^, with STEAP4^pos^ showing significant reductions relative to the stromal compartment in CeD samples in contrast to observations in inflamed ileum and colon from Crohn’s disease (Fig. 5b,c). As previously mentioned, relative to the total sample, DES^pos^ ACTA2^pos^ SMC showed significant decreases in CeD biopsies (Fig. 5c).

**Figure 5.**
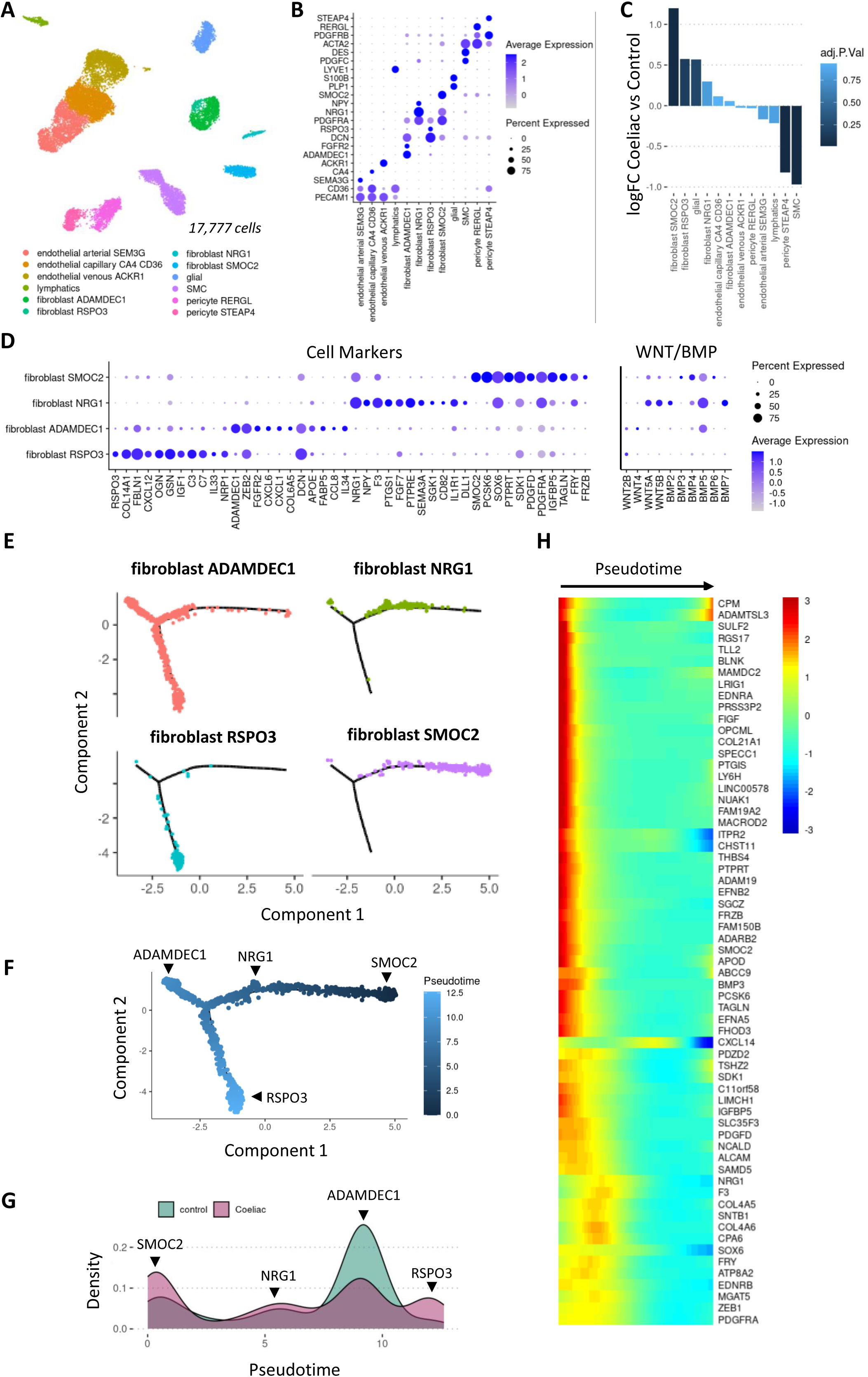
Coeliac Disease is characterized by unique tissue relevant stromal cellular subsets reflective of disease re-modeling and repair. A) UMAP visualization of stromal compartment. B) Expression of gene markers per cell type annotation. C) Differential abundance analysis of stromal compartment showing logFC of cell types in Coeliac vs control samples. D) Expression of markers for fibroblast subtypes (left) and expression profiles of WNT/BMP genes (right). E) Pseudotime trajectories used to distinguish fibroblast subtypes, visualized by cell type annotation along trajectory. F, G) Pseudotime visualization of trajectories starting at the SMOC2 fibroblast branch with associated density plot of pseudotime per condition. H) Heatmap of differentially expressed genes that increase along the NRG1/SMOC2 trajectory.

Four distinct subpopulations of fibroblasts were characterized across control and CeD samples annotated as RSPO3^pos^, ADAMDEC1^pos^, SMOC2^pos^ and NRG1^pos^ fibroblasts (Fig. 5a). SMOC2^pos^ fibroblasts significantly increased in number relative to the stromal compartment in CeD, in addition to increases in RSPO3 and NRG1 subsets (Fig. 5b,c). RSPO3, which encodes R-spondin 3, a WNT signaling progenitor, was uniquely expressed on a distinct fibroblast subtype in addition to OGN, GSN, C3 and IL33, hallmark features of this crypt-supporting fibroblast. ADAMDEC1 fibroblasts minimally changed in number in CeD, but showed high levels of ADAMDEC1, FGFR2, chemokines CXCL1 and CXCL6, and IL34 among others, suggesting a role in signaling to local immune cell types. ADAMDEC1, previously characterized in the context of UC and DSS colitis, prevents the aberrant accumulation and disorganization of ECM and thus has a protective role in cellular homeostasis in the large intestine^39, 40^, potentially highlighting a disorganized ECM turnover in CeD. Notably, we describe two distinct clusters of PDGFRA^high^ fibroblasts comprising the NRG1^pos^ and SMOC2^pos^ subsets, in contrast to the PDGFRA^low^, RSPO3^pos^ and ADAMDEC1^pos^ fibroblasts. Both populations share expression in PDGFRA, NRG1, F3, SOX6, and FRY to various degrees; NRG1^pos^ fibroblasts are distinctly characterized by high expression of NPY and DLL1 among others, whereas SMOC2 ^pos^ fibroblasts are characterized by high expression of SMOC2, PCSK6, PTPRP among others (Fig. 5d). PDGFRA^high^ fibroblasts have previously described characteristic of subepithelial fibroblasts in the colonic mucosa which orchestrate the differentiation of intestinal epithelial cells and have a role in villus formation, whereas RSPO3 fibroblasts (PDGFRA^low^) support crypt epithelial functions and ADAMDEC1 fibroblasts have been located around crypts and in lamina propria^41^. This is thought to be achieved in part through maintenance of a gradient of WNT and BMP mediated signaling along the crypt villus axis ^42, 43, 44, 45^, reflected by the canonical WNT2B expression in RSPO3 and ADAMDEC1 fibroblasts and non-canonical WNT5A, WNT5B and BMP expression profiles of SMOC2^pos^ and NRG1^pos^ fibroblasts (Fig. 5d, right). Interestingly, within the PDGFRA^high^ fibroblasts, BMP6 is distinct to SMOC2 fibroblasts and BMP7 to NRG1 fibroblasts. Overall, these data provide a novel view into two clearly distinct subsets within PDGRA^high^ fibroblast populations that suggest a more nuanced role of subepithelial fibroblasts within the duodenum.

To further contextualize the CeD-enriched PDGRA^high^ NRG1 and SMOC2 fibroblasts, yet given the limited cell numbers per fibroblast cluster, we sought to leverage pseudotime analysis to highlight transcriptional differences between these fibroblast populations in CeD. Pseudotime analysis (Fig. 5e) resulted in a separate SMOC2 branch that included NRG1 fibroblasts along the trajectory, distinct from both ADAMDEC1 and RSPO3 fibroblast branches. CeD samples showed higher densities of cells along SMOC2 and RSPO3 branches (Fig. 5f,g), in line with increases seen with differential abundance analysis. Differential gene expression along SMOC2 trajectory pseudotime was characterized by increases in genes previously described, including SMOC2, NRG1, PDGFRA, F3, SOX6, as well as COL4A5 basement membrane and BMP3 (Fig 5h). This is consistent with previous studies on PDGFRA^pos^ subepithelial fibroblasts that support the villus structure and epithelial cell differentiation. Combined with the increase in abundance of these SMOC2 branch cells (NRG1 and SMOC2 fibroblasts) in CeD, these fibroblasts represent distinct stromal changes associated with epithelial differentiation and crypt stem cell lineages and may reflect a key repair pathway significantly upregulated in CeD.

### Myeloid derived IL1β signal to NRG1^pos^ fibroblasts modulating duodenal epithelial cell states in disease

Highlighting the relevance of the SMOC2 branch fibroblasts in CeD, many of the genes that increased along the SMOC2 pseudotime branch, expressed across NRG1 and SMOC2 fibroblasts, also overlapped with genes that were significantly upregulated in the fibroblast compartment of CeD samples (Supplementary Table 1).

To analyze the upstream causal factors of these differential changes in fibroblasts, we performed Nichenet cell-cell communication analysis on the differentially increased fibroblast genes as target genes within the SMOC2 branch fibroblasts (i.e., SMCO2 fibroblasts and NRG1 fibroblasts). Nichenet ligand activity analysis revealed several ligands with regulatory potential on these downstream fibroblast target genes (predominantly expressed in SMCO2 fibroblasts and NRG1 fibroblasts) through their corresponding putative receptors (Fig. 6a, 6b, Extended Fig. 3). These ligands included key fibroblast activating mediators such as IL-1β, TGF-β1, and EDN1 (Fig. 6a). Notably, top ranked ligands also included IFN-γ (Fig. 6a), expressed in the T/NK compartment (Fig. 6b) consistent with the previously discussed CellChat IFN-γ signaling analysis from effector T cells to IFN—γ receptor expressing SMOC2^pos^ fibroblasts and NRG1^pos^ fibroblasts (Fig. 2f).

**Figure 6.**
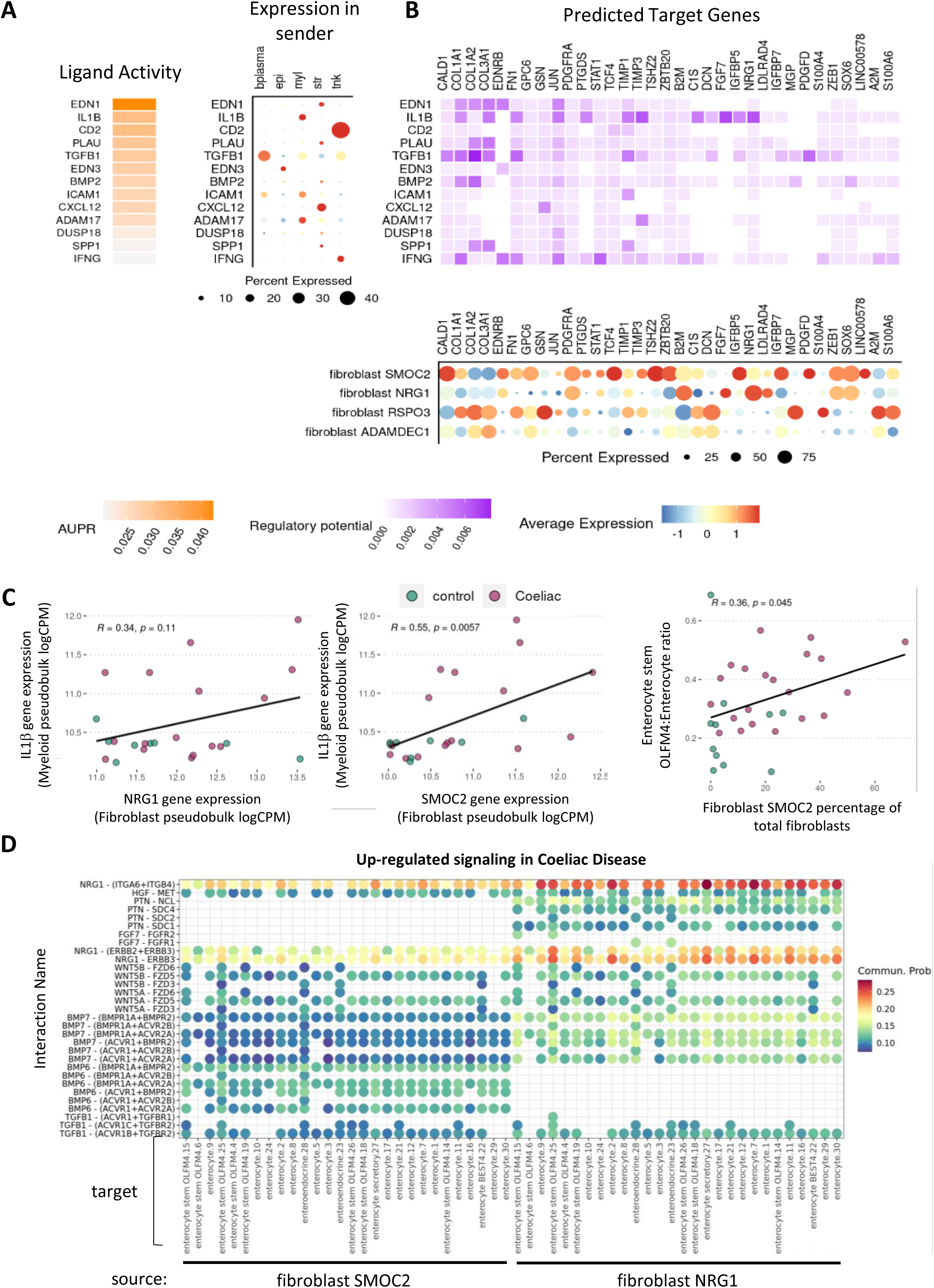
Myeloid-derived IL1β upstream of Coeliac transcriptomic profile of NRG1-expressing fibroblasts, supporting epithelial differentiation signaling in disease. A, B) NicheNet ligand activity analysis using significantly increased genes within fibroblast compartment in Coeliac samples analyzed as downstream target genes in NRG1 and SMOC2 fibroblasts. A) Area under the precision-recall curve (AUPR) was used to rank ligand predicted activity based on observed target gene expression with dotplot visualizations of expression of ligands across compartments and B) expression of targets across fibroblast subtypes and heatmap of regulatory potential between ligands and target genes. C) Left, Correlation of IL1B gene expression from myeloid compartment pseudobulk with NRG1 and SMOC2 gene expression from fibroblast pseudobulk per sample; R indicates Pearson correlation and associated p-value. Right, Correlation of enterocyte stem OLFM4 to enterocyte ratio with SMOC2 fibroblasts as percentage of total fibroblasts; R indicates Pearson correlation and associated p-value. D) CellChat bubble plots showing significantly up-regulated secreted signaling between NRG1 and SMOC2 fibroblasts and epithelial cell clusters (marked by annotation with Leiden cluster) in Coeliac vs control samples.

In particular, the IL-1β ligand, expressed in the myeloid compartment (Fig. 6a), associated with CeD pathogenesis, was predicted to have high regulatory potential on NRG1 (Fig. 6b), a gene increased along the SMOC2 trajectory with shared expression in SMOC2 and NRG1 fibroblasts. To substantiate this inferred biological association, subject-specific myeloid compartment gene expression was correlated to the paired subject-specific fibroblast compartment gene expression. Myeloid IL1B gene expression correlated with both NRG1 gene expression and SMOC2 gene expression in fibroblasts across samples, with highest expression in CeD samples (Fig. 6c). NRG1, a marker of this SMOC2 branch, has been reported to induce fetal-like reprogramming of the intestinal epithelium and supports epithelial differentiation^46^. SMOC2^pos^ fibroblast percentage significantly correlated with the enterocyte stem:enterocyte ratio across samples (Fig. 6c), highlighting these relationships between stromal support cells and epithelium in CeD. In support of biological association of IFNG ligand signaling to fibroblasts, subject-specific cell compartment correlations also showed that IFNG expression in T cells correlated with NRG1 and SMOC2 gene expression in fibroblasts across samples (Extended Fig. 4a). Furthermore, IFNG expression in T cells also correlated with IL1B myeloid expression, consistent with the previously discussed CellChat IFN-γ signaling analysis from effector T cells to IFNG receptor-expressing myeloid populations (Extended Fig. 4b). These data-driven associations with IFNG highlight potential mechanisms of downstream cross-compartmental effects of the effector T cell population in CeD in affecting both stromal and myeloid populations within the tissue.

Given their putative role in supporting epithelial differentiation, we next sough to analyze the downstream signaling of these SMOC2 branch fibroblasts to the epithelium in CeD. CellChat analysis of the secreted signaling in SMOC2 branch fibroblasts revealed several interactions with epithelial cells that were significantly upregulated in CeD that included NRG1 itself, in addition to HGF, PTN, FGF7, TGFB1, BMP6, BMP7, WNT5A and WNT5B ligands, known to support epithelial barrier formation and differentiation (Fig. 6d). These results are supported by previous work demonstrating the direct binding of NRG1 to integrins αvβ3 and α6β4 thereby mediating NRG/ErbB signaling in fibroblast cell types, highlighting the increased role of EGF family signaling to regulate duodenal epithelium in CeD ^47, 48^(Fig. 6d).

In totality, the combined differential changes in CeD in cell abundance, gene expression and cell-cell interactions across cellular compartments propose an interesting tissue cellular cross talk between immune and non-immune subsets unique to active CeD. Here, myeloid derived IL-1β, increased in CeD, mediates signaling to PDGFRA^high^ SMOC2 and NRG1 fibroblast subsets. These fibroblasts then secrete ligands, such as WNT, BMP and NRG1, known to support intestinal epithelium. Receptor-ligand interactions revealed significant communication between these fibroblast subsets and the enterocyte populations, suggesting the role of these fibroblasts to support the epithelial reprogramming of the increased stem/crypt epithelial fraction (i.e. crypt hyperplasia) in CeD (Illustration 1; Proposed schematic for cellular interactions in CeD). This data proposes a key role related to a previously undescribed myeloid-stromal-enterocyte interface modulation in CeD pathogenesis.

## Discussion

Coeliac Disease remains a challenging condition, with significant unmet clinical need for those living with the disorder, particularly those individuals with refractory disease ^15^. Despite its global prevalence, no therapeutic approaches have been approved for the treatment of CeD, with recent unsuccessful studies targeting gluten degradation and other immune tolerance mechanisms^49^. To this end, identification of novel disease relevant cellular dynamics and interactions, associated with CeD pathogenesis, are required to characterize tissue specific mechanisms responsible for disease pathogenesis, repair, and resolution. Critical to achieving this aim is the characterization of non-immunological cell types associated with tissue inflammation, remodeling, and homeostasis.

This study provides the first comprehensive single-cell transcriptomics evaluation of CeD, characterizing both immune and non-immune cell populations. Additionally, integration of cellular interaction analysis proposes a tissue-level context of how individual cellular subsets interact to contribute to inflamed mucosa dynamics of CeD individuals with gluten exposure. Our dataset provides novel human information on the enterocyte, myeloid and stromal compartments in CeD, and enables the characterization of cell type specific differences in disease. Further, our data points to a specific cellular interaction between myeloid, stromal and enterocyte compartments contributing to cellular dysfunction in CeD.

These data contribute several new key findings for CeD. First, the major cellular differences between control and inflamed CeD duodenal tissue are described. Disease-associated cell types, including Treg, effector CD8^+^ T, γδ T cells and T_fh_-like CD4^+^ T cells were significantly enriched in CeD, consistent with previous findings by Atlasy et al ^35^. In addition to changes in cellular frequency, distinct T cell transcriptional states were associated with CeD, including type 1 effector biology, broad activation markers and pathways associated with cell stress. These cellular subsets mediated cell-cell interactions via a variety of downstream ligands, including IFN-γ and IL21, which were exclusively expressed in CeD. This gluten induced cytokine changes directly interacted with a variety of downstream cell types including myeloid and stromal subtypes, with broad immune modulation capability.

Second, in addition to T and B cell changes, we identified several myeloid cellular subsets enriched in CeD. This is highly relevant, as to date, limited myeloid transcriptomic data is available in CeD, and it was unclear if myeloid subpopulations were selectively enriched or altered in CeD. Here, we found increases in DC1, DC2 and monocytic cellular subsets. Interestingly, these subsets had an enhanced cellular activation phenotype, characterized by IL-1β, FOS and TGF-β1 co-expression patterns. In addition to pro-inflammatory cytokines, CeD specific myeloid cells also expressed significant amounts of CCL4, CCL5, CXCL8 and CXCL16 communicating with both endothelial, stromal and T/B cell types. Furthermore, tissue resident macrophage subsets, associated with intestinal immune tolerance ^50^, were significantly decreased relative to the myeloid compartment in CeD, demonstrating a cellular shift within CeD tissue towards inflammatory myeloid populations.

Third, our work demonstrates a significant shift in enterocyte frequencies, phenotypes, and transcriptional states within the mucosa of CeD patients. Here, CeD samples had a significant enrichment in OLFM4 stem^51^ and secretory enterocytes, whereas absorptive enterocytes were reduced. Moreover, enterocyte transcriptional module analysis demonstrated a shift towards a progenitor state with reduced absorptive capacity, as highlighted by the stem:enterocyte ratio and were associated with clinical histological parameters, providing single cell data evidence of the hallmark CeD epithelial histologic features of villous atrophy and crypt hyperplasia.

Fourth, our work identifies unique stromal immune cellular subsets associated with CeD pathogenesis. These fibroblasts characterized as NRG1^POS^, SMOC2^POS^ and RSPO3^POS^ were enriched in the mucosa of patients with CeD. These subsets have been implicated in the development and homeostasis of epithelial cells within the intestinal niche. Our data reveal a previously undefined shift in PDGFRA^low^ fibroblasts (i.e., RSPO3^pos^) in CeD, and increases in two similar yet distinct PDGFRA^high^ fibroblasts, NRG1 and SMOC2 fibroblasts, thus contributing to the underlying tissue biology of CeD.

Previous reports have noted the role of intestinal fibroblasts in maintaining the homeostatic stem cell niche as well as supporting differentiation and maintenance of enterocytes along the crypt-villous axis. In human development, villous fibroblast progenitors are PDGFRA^high^ and express the EGF-family ligand NRG1, which provide essential epithelial crosstalk to regulate differentiation, morphogen gradients (e.g., BMP), and villus structure formation. Administration of NRG1 to adult murine epithelium and human epithelial organoids has been shown to affect secretory lineage differentiation and increase cellular diversity ^48, 52^. Similarly, Secreted modular calcium-binding protein 2, SMOC2, has previously been postulated to facilitate intestinal crypt formation through inhibition of BMP signaling, thereby regulating the BMP morphogen gradient in villi ^53, 54^. Our data has revealed an increase in fibroblast subsets expressing NRG1, both SMOC2 and NRG1 subsets, associated with CeD. This enhanced signaling may differentially impact the enterocyte lineage, as reflected by the increase in stem and secretory enterocytes observed in our study. In addition to PDGFRA^high^ SMOC2 and NRG1 fibroblasts, RSPO3 expressing fibroblasts have also been shown to be differentially modulated following injury, resulting in increased epithelial regeneration. Both RSPO3 and PDGFRA^high^ fibroblasts likely increase interactions with the hyperplastic stem/crypt epithelium to help restore the tissue to its homeostatic state. Interestingly, inflammatory activated fibroblasts identified in the large intestine of IBD patients^21, 22^ and characterized as IL11^pos^IL13Ra2^pos^ were absent in the duodenal mucosa of CeD, reflecting potential differences between the fibroblast niche associated with inflammation in distinct anatomical compartments within the intestine. Taken together, these data revealed a shift in small intestine supporting fibroblast subsets in CeD, reflecting dynamic stromal cellular interactions to support epithelial reprogramming and repair mechanisms in the diseased CeD mucosa.

Further, our data has also revealed a decrease in ADAMDEC1 expression in fibroblasts in CeD, these cells have been implicated in fibroblastic response to tissue injury and healing in IBD, displaying a proliferative phenotype and are a major source of collagen^39^. Therefore, defective healing, in the absence of ADAMDEC1^pos^ cells, may contribute to an impairment in histologic healing in CeD. Additional analysis is required to evaluate the mechanistic contribution of this cell type in the small intestine during health and homeostasis.

Based on their significant cellular contribution and dysregulation in CeD, myeloid, stromal and enterocyte subsets were evaluated to interrogate specific cellular communications between these subsets. Our cell chat analysis revealed substantial cross talk between T, myeloid, stromal and enterocytes in disease. Analysis of upregulated genes in CeD fibroblasts indicated high regulatory potential of ligands coming from CeD monocytic lineages (including IL-1β). Previous reports from Friedrich et al., have highlighted the role of IL-1β in IBD and the impact on fibroblast disease states. Here, the authors found that activated fibroblasts in the ulcer bed had significant IL1R, but not TNF, dependence. Furthermore, these IL1R^pos^ fibroblasts were associated with refractory disease in UC and CD ^55^. Additionally, it has recently been demonstrated that stromal IL-1R1 activity can support epithelial regeneration in the mouse intestine ^56^. With this in mind, we evaluated the predicted target genes associated with the unique fibroblast subsets represented in our CeD samples. Interestingly, IL-1β was identified as a top gene associated with SMOC2^pos^ and NRG1^pos^ populations suggesting cellular cross talk between the infiltrating myeloid and unique duodenal stromal populations. NRG1 has been reported to impact the fetal-like reprogramming states of the intestinal epithelium and support epithelial differentiation, consistent with the CeD associated enterocyte phenotypes described in our study to be increased in CeD. Furthermore, analysis of the downstream secreted signaling from these SMC2^pos^ and NRG1^pos^ cells revealed significant interactions between various epithelial cells, mediated by NRG1 and other BMP and WNT ligands.

As such, one could speculate that myeloid derived IL-1β, as a consequence of gluten specific IFNγ signaling is a key regulator of downstream cellular crosstalk. In support of the above, we confirmed significant cellular cross communication between CeD induced CD8^POS^ T cells and monocytic cell lineages. These monocytic lineages also express high levels of the IFNR1 and IFNR2. Myeloid derived IL-1β results in modulation of the NRG1^POS^ and SMOC2^pos^ fibroblasts resulting in a shift in the enterocyte stem: mature enterocyte ratio, highlighting the relationship between stromal cells and the epithelium in Coeliac disease (Illustration 1).

Taken together, our study of control and active CeD, based on high-resolution scRNAseq transcriptomics approaches, has revealed a previously undescribed disease associated stromal-enterocyte-immune interaction. These data provide a deeper understanding of the cellular networks underlying the mucosal inflammation in disease while informing on new possible therapies targeting both the myeloid and stromal compartment as an effect treatment in CeD.

## Material and Methods

### Patient selection

The SUCCEEDS Study (*S*cience *U*nderlying *C*oelia*C E*volution: *E*xplanations, *D*iscoveries, and *S*olutions) is a prospective study of paediatric coeli disease (age under 17 years), based at Children’s Health Ireland, Dublin, approved by the local Research Ethics Committee (REC-104-22). Clinical, laboratory, endoscopic and histological data are collected on bespoke case report forms (Supplementary Table 1). Samples were obtained from participants at the time of diagnostic endoscopy, following fully informed consent. All participants were consuming a full gluten-containing diet at the time of endoscopy. Cases were identified based on subsequent histological confirmation and Marsh scoring. Normal duodenal histology was confirmed in all control subjects.

### Sample collection and cryopreservation

Duodenal biopsies collected from participants were placed immediately in cryovials containing 1ml of cryostat C10 cell cryopreservation media (#C2874, Sigma Aldrich) on ice and were placed in -80 within 2 hours of collection using a coolcell container.

### Dissociation and live cell enrichment of cryopreserved duodenal biopsies

Cryopreserved biopsies were thawed by immersing in a 37°C water bath for ∼ 2 min with periodic shaking. RPMI with 10% FCS was added to each biopsy, dropwise in a 50 ml tube. The biopsies were washed in serum free RPMI over a cell strainer. Single cell suspensions were made by placing biopsies in a mixture of Liberase TM (SIGMA – 5401127001) and DNase (SIGMA - D5025-15 KU) while slowly rotating at 37°C (*Smilie et al*.,). Dissociated biopsies were washed with 0.1% BSA over a cell strainer. Dead cells were removed using EasySep™ Dead Cell Removal (Annexin V) Kit (Stemcell Technologies - 17899). Cells counts were determined using AOPI staining solution and a Nexcelom fluorescent cell counter.

### Single cell processing

Single cell FASTqs were processed using cellranger v7.0 followed by cellbender ambient RNA removal [Fleming 2023]. Seurat v5 package in R was used for single cell processing and analysis. DoubletFinder v2.0.4 was used per sample to score and identify doublets. For quality control (QC), cells with cellbender-derived cell probably < 0.5, nCount < 200, nFeature < 200, percent mitochondrial reads (percent.mt) > 50, were removed. SCTransform was used to normalize, scale, and for variable feature selection. Total cell object was integrated between samples using Harmony and clustered into broad compartments (str – stromal, imm – immune, and epi – epithelial) for compartment-specific QC where doublets and low quality cells/clusters were filtered out. The imm compartment was reclustered and divided into TNK (T cell, ILC, NK cells), bplasma (B cells and plasma cells) and myl (macrophage, DC, monocyte, Mast). Given the higher occurrence of percent.mt in epithelial cells, the epi compartment was refiltered with a more inclusive filter of percent.mt < 40, while the rest were refiltered to percent.mt < 20. Leiden clusters were then used for cluster-based annotations according to data-driven cluster-specific signals alongside reported cell markers.

### Single Cell Library Preparation

scRNA libraries were prepared using the Chromium Next GEM Single Cell 3ʹ Kit v3.1 (10X Genomics) according to manufacturer’s instruction on the Chromium Controller (10X Genomics). Sequence-ready libraries were validated and quantitated using the Agilent High Sensitivity D1000 ScreenTape Kit on the TapeStation 4200 System (Agilent Technologies). Individual libraries were normalized to 3nM and quantified by qPCR using the KAPA Library Quantification Kit – Illumina/ROX Low (Roche) on the QuantStudio 12K Flex Real-Time PCR System (Applied Biosystems) according to manufacturer’s instructions. The libraries were then pooled to 3nM and quantified again by qPCR as described above. The quantified library pool was denatured and diluted according to the NovaSeq 6000 Denature and Dilute Libraries Guide and loaded onto a NovaSeq 6000 System (Illumina) for paired end sequencing. BCL conversion and demultiplexing was performed using the bcl2fastq2 script (v2.20, Illumina).

### Differential abundance analysis

To assess changes in cell type composition, we address the compositionality of the cell abundance count data by using a compositional transformation, in particular the centered log-ratio (CLR) transformation [Aitchison, 1982]. Upon transformation, the counts are used as a response variable in a linear model that is fitted for each cell type separately. For linear model fitting, we leverage the limma framework [Smyth, 2004], additionally allowing us to account for the heteroscedastic nature of the counts and adopting empirical Bayes shrinkage on the linear model residual variances.

For statistical inference, we acknowledge the recent work of Zhou et al. 2022, showing that log fold-changes are biased with respect to the true effect sizes on the cell type’s absolute abundances, and thus use the bias correction approach as suggested in Zhou et al. (2022). Statistical inference is assessed using moderated t-tests [Smyth, 2004] and p-values are adjusted according to the Benjamini-Hochberg false discovery rate [Benjamini & Hochberg, 1995].

### Differential gene expression analysis

Differentially expressed (DE) genes were analyzed after pseudobulk mean aggregation per cell type or cell group of interest using the muscat package for aggregation. Subject samples containing less than 20 cells were removed prior to analysis. Immunoglobulin genes (except in Bcell/plasma cell contrasts), mitochondrial genes, and genes with less than 10 counts across all samples were filtered out. The edgeR package was used to calculate normalize factors and limma-voom analysis was used for DE. DE gene summaries were defined as p-value < 0.05.

### Gene module construction and differential gene module analysis

Prior to gene correlation networks, the metacell package was used to construct metacells per compartment (str, epi, TNK, bplasma, myl). Briefly, harmony embeddings were supplied to the metacell algorithm prior to the balanced K-nearest neighbors graph construction step. The K value was set to 200 for each compartment, except for TNK, which was set to K = 400 to account for the lower average UMI counts in the TNK compartment. Metacell pseudobulk object was constructed by summing the SCT counts of each cell per metacell and then processed and normalized using edgeR where genes less than 10 counts in all samples were removed. Gene correlation networks were constructed using graphia based on Pearson correlation where correlation thresholds optimized visually per compartment. Markov clustering of genes (MCL) was used to identify and define correlated/co-expressed gene modules for DE module analysis. Modules were identified as the MCL cluster number with either a gene contained in the module or according to a consensus cell type it is known to represent (e.g., tnk7.Treg module contains well known FOXP3 Treg gene). The Supplemental table X contains a list of all modules per compartment. Gene set variation analysis (GSVA) was then used to score modules per cell type/contrast of interested on pseudobulk objects constructed previously and differential analysis was performed using the limma package.

### Cellular interaction analysis

CellChat V2 was used to infer intercellular communication across all populations and to identify differential signaling between Coeliac and healthy samples. CellChat quantifies communication between two cell groups by calculating an interaction score based on the average expression values of a known ligand and its receptor, as well as their cofactors, using the law of mass action. Ligand-receptor pairs that were expressed in at least 5% of cells and present in a minimum of 10 cells per cluster were selected^57, 58^.

The NicheNet R package was used to analyze upstream ligand activity predictive of downstream target gene expression. The fibroblast pseudobulk DE genes were identified as the gene sets of interest evaluated in the receiver target population of fibroblast SMOC2 and fibroblast NRG1 cells (cells on the SMOC2 pseudotime trajectory branch). All cells were used as putative sender cells. Top ranked ligands were identified according to the ligand activity analysis workflow, where receiver and sender expression was filtered to be expressed in at least 5% of cells and the top 2000 targets per ligand that were considered as potential “active ligand-target links.” In line with NicheNet validation studies, area under the precision-recall curve (AUPR) was used to rank ligand predicted activity with observed target gene expression.

### Pseudotime analysis

Pseudotime analysis of the fibroblasts was performed using monocle. Immunoglobulin, ribosomal, and mitochondrial genes were removed prior to analysis. To construct the pseudotime trajectories in the context of coeliac disease understanding, genes were first ordered according to the q-value from on Coeliac versus control differential analysis using monocle’s differentialGeneTest function. To identify genes that were differentially increased along the SMOC2 trajectory, the pseudotime State at the end of the SMOC2 trajectory was set as the root state prior to pseudotime differential gene analysis. Genes with a pseudotime DE q-value < 1e-50 were then used for heatmap visualization.

## Data Statement

The dataset generated during the current study are publicly available on https://www.ncbi.nlm.nih.gov/ (Accession number: Pending). All analytical methods, including statistical codes, software information, and algorithms used in this study, will be made available upon request to the corresponding authors. Further information and requests for resources and reagents should be directed to and will be fulfilled by the lead contact

## Author Contributions

DR, KS, CB, DH, JR, LT, KP, HL, SH, DR, PW performed the experiments and analyzed the data. DR, KS, BR, SH, DR, PW contributed to the writing of the manuscript. DR, PW and SH provided research funding, research supervision or research materials. DR, PW, SH conceived the research, led the project, analyzed the data, interpreted the results, and wrote the manuscript.

## Acknowledgements

This work has been supported by Janssen Research & Development, LLC, a wholly owned subsidiary of Johnson & Johnson. Additional funding by Science Foundation Ireland (21/FFP-P/10135) and Childrens Health Foundation to PTW and SH (RSFG-21-ACC07). We acknowledge the input of Drs Michael McDermott, Maureen O’Sullivan and John O’Neill as the reviewing pathologists, and Roberta Hack Mendes and Anna Dominik who supported patient recruitment. We also thank Drs Jan Wehkamp, Weiwei Schultz, Esi Lamousé-Smith, Carolyn Cuff, Arun Kannan, Tae Lee and Tom Freeman for helpful discussion and feedback.

---

**Diagram 1:**
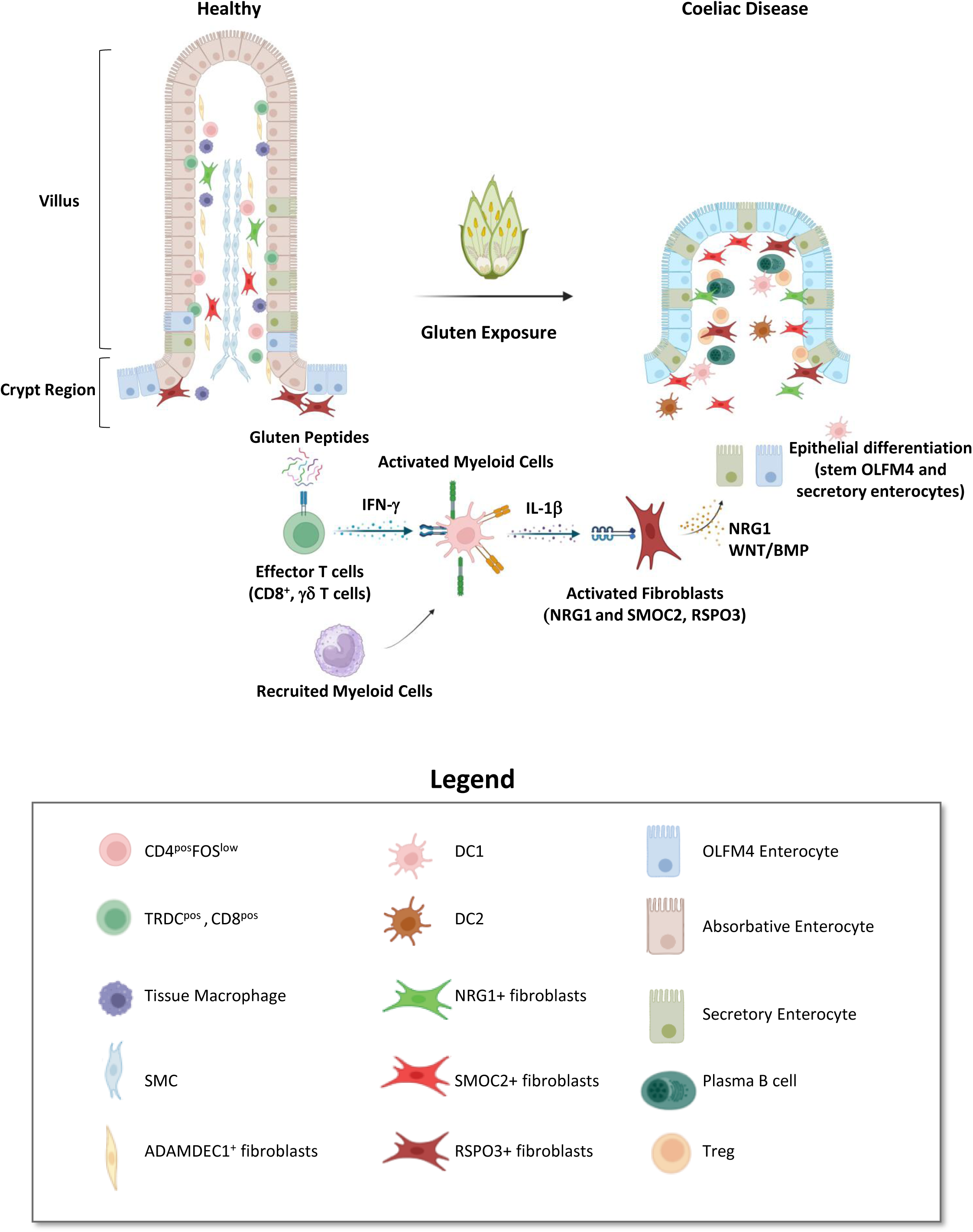
Coeliac Disease Pathogenesis. A) Schematic representation of the healthy duodenal mucosal. Homeostasis is characterized by significant levels of absorbative enterocytes, smooth muscle cells, tissue resident macrophages and a diverse array of lymphoid cellular subsets, including naïve, IELs and TRDC+TYROBP+ CD8+ T cells. Upon gluten exposure in a Coeliac Disease individual, a significant cellular shift is noted within the mucosa. With significant increases in B cells, monocyte/myeloid populations, and fibroblast subsets. Further, increases in T_regs_ and T_fh_ also track with disease. Increases in OLFM4 (stem-like) enterocytes, and secretory enterocytes are also a hallmark of disease, with significant shifts notes in the enterocyte OLFM4: Enterocyte ratio compared to homeostasis. NRG1, RSPO3 and SMOC2 fibroblasts are also increased in disease. **A model for T-myeloid-stromal-epithelial crosstalk in Coeliac Disease.** B) Gluten exposure results in the rapid activation of effector T cells, including CD8+, δγ T cells, as characterized by the release of IFN-γ, resulting in signaling to monocytic and DC2s, expressing IFNR1/2. IL-1β release from activated myeloid subsets in response to upstream IFN-g, results in cellular activation of tissue resident cells including NRG1+ and SMOC2+ fibroblasts. Activation of duodenal fibroblasts, results in enterocyte remodeling and repair, measured by NRG1 and WNT/BMP ligand expression. WNT/BMP/NRG1 expression results in the differentiation of enterocytes along a stem OLFM4 and secretory enterocyte lineage observed in disease samples.

