## Supplementary Figures 1-3 for "Immune signaling mediates stromal changes to support epithelial reprogramming in Celiac duodenum"

**A**

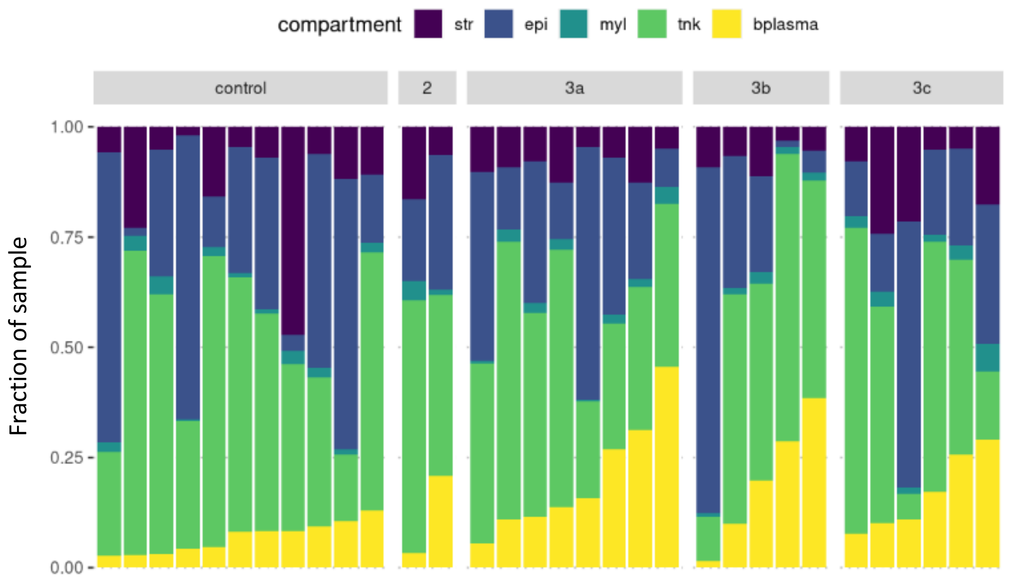


**Supplemental Figure 1. Single-cell atlas of duodenal biopsies reveals a shift in cell type composition with Marsh score.** A) UMAP visualization and histogram of all cells collected, ordered by increasing plasma cell fraction, characteristic of Coeliac disease tissue, stratified based on Marsh scores

A

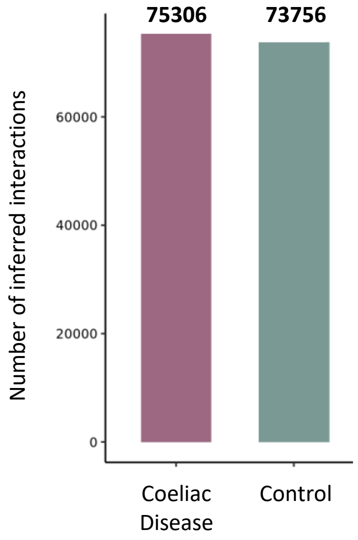


**Supplemental Figure 2. CellChat comparison of Coeliac Disease and Control signaling** A) Bar chart outlining the total number of interactions in Coeliac and Control samples.

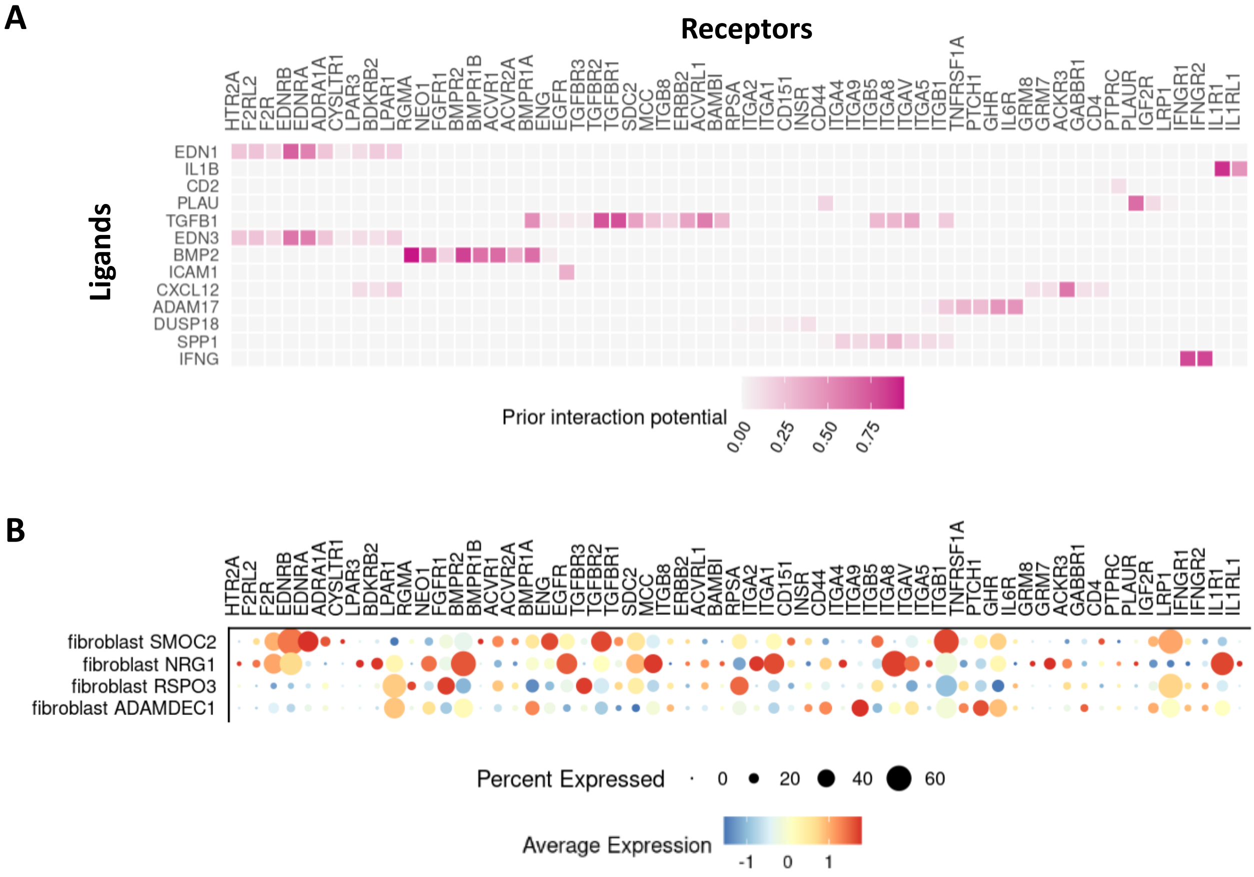


**Supplemental Figure 3. Ligand-receptor interaction upstream of Coeliac transcriptomic profile of NRG1-expressing fibroblasts.** A) Heatmap of NicheNet's predicted ligand-receptor interaction potential supporting ligand activity analysis, displaying receptor genes (expressed in NRG1 and SMOC2 fibroblasts) related to top ranked ligands. B) Dotplot showing expression of receptor genes across all fibroblast subsets.

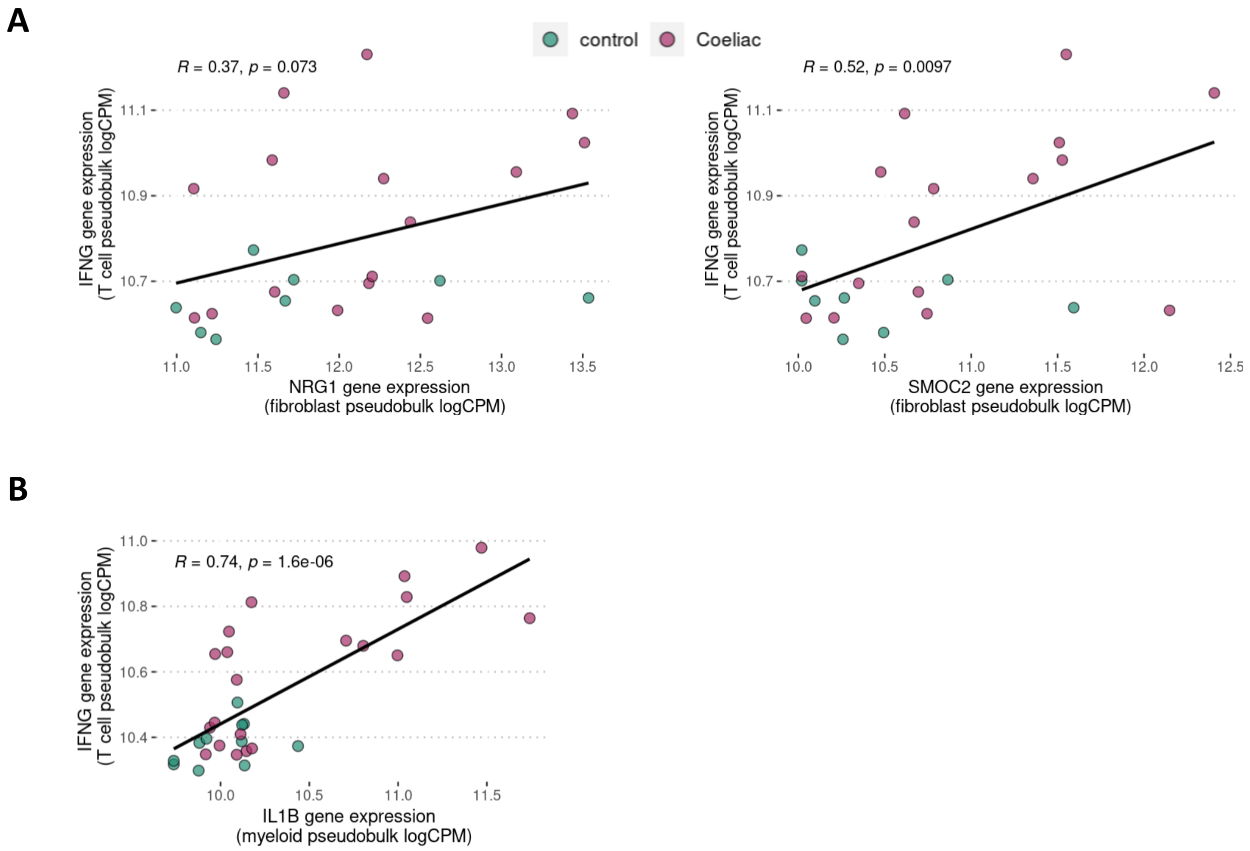


**Supplemental Figure 4. T cell-derived IFNG is correlated to CeD-associated fibroblast genes, NRG1 and SMOC2, and CeD-associated myeloid gene, IL1B.** A) Correlation of IFNG gene expression from T cell (excluding NK, ILC) compartment pseudobulk with NRG1 and SMOC2 gene expression from fibroblast pseudobulk per sample; R indicates Pearson correlation and associated p-value. B) Correlation of IFNG gene expression from T cell (excluding NK, ILC) compartment pseudobulk with IL1B gene expression from myeloid compartment pseudobulk; R indicates Pearson correlation and associated p-value.
