## Supplementary Table 1 for "Immune signaling mediates stromal changes to support epithelial reprogramming in Celiac duodenum"

**Supplemental Table 1:**

| | Control (n=11) | Coeliac (n=21) | Significance |
|---|---|---|---|
| Sex (Male/Female) | Male - 5<br>Female - 6 | Male - 6<br>Female - 15 | |
| Age (Years) | 3.5 - 17.0 | 4.1 - 15.9 | |
| tTG Absolute (mean +/− SD (range)) | 12.10 +/− 19.288 (0.0-59.0) | 48.76+/−42.136 (0.0-128.0) | * |
| [1.] tTG-rel-ULM (mean+/− SD (range)) | 1.718 +/− 2.752 (0.0-8.4) | 6.952+/−6.175 (0.0-18.3) | * |
| [2.] Marsh 1 | 0 | 0 | |
| Marsh 2 | 0 | 2 | |
| Marsh 3a | 0 | 8 | |
| Marsh 3b | 0 | 5 | |
| Marsh 3c | 0 | 6 | |

**Table 1: Patient details for samples included in scRNAseq analysis.**

tTG – tissue transglutaminase; SD standard deviation.

[1] tTG values expressed relative to the upper range of normal of the test assay.

[2] Based on the location of the most severe histopathology finding in each patient.

Statistical analysis performed by unpaired student’s t test (* p≥0.05)
